# Mapping of TNTs using Correlative Cryo-Electron Microscopy Reveals a Novel Structure

**DOI:** 10.1101/342469

**Authors:** Anna Sartori-Rupp, Diégo Cordero Cervantes, Anna Pepe, Elise Delage, Karine Gousset, Simon Corroyer-Dulmont, Christine Schmitt, Jacomina Krijnse-Locker, Chiara Zurzolo

## Abstract

The harmonious orchestration of intercellular communication is essential for multicellular organisms. One mechanism by which cells communicate is through long, actin-rich membranous protrusions, called tunneling nanotubes, that allow for the intercellular transport of various cargoes, including viruses, organelles, and proteins between the cytoplasm of distant cells *in vitro* and *in vivo*. Over the last decade, studies have focused on their functional role but information regarding their structure and the differences with other cellular protrusions such as filopodia, is still lacking. Here, we report the structural characterization of tunneling nanotubes using correlative light- and cryo-electron microscopy approaches. We demonstrate their structural identity compared to filopodia by showing that they are comprised of a bundle of functional individual Tunneling Nanotubes containing membrane-bound compartments and allowing organelle transfer.

## Introduction

Tunneling nanotubes (TNTs) have been defined as long, thin, non-adherent membranous structures that form contiguous cytoplasmic bridges between cells over long and short distances ranging from several hundred nm up to 100 µm ^1^^-^^4^. Over the last decade, scientific research has effectively improved our understanding of these novel structures and underscored their role in cell-to-cell communication, facilitating the bi- and uni-directional transfer of compounds between cells, including: organelles, pathogens, ions, genetic material, and misfolded proteins ^5^. Altogether, *in vitro* and *in vivo* evidence has shown that TNTs can be involved in many different processes such as stem cell differentiation, tissue regeneration, neurodegenerative disorders, immune response, and cancer ^2,6-11^.

Although these *in vitro* and *in vivo* studies have been informative, the structural complexity of TNTs remains largely unknown. As a result, TNTs have been regarded with skepticism by one part of the scientific community ^5,12^. Two major issues that remain to be clarified are whether these protrusions are different from other previously studied cellular processes such as filopodia ^13^ and whether their function in allowing the exchange of cargos between distant cells is due to direct communication between the cytoplasm of distant cells or to a classic exo-endocytosis process or a trogocytosis event ^14^.

Addressing these questions has been difficult due to considerable technical challenges in preserving the ultrastructure of TNTs for electron microscopy (EM) studies. To date, the ultrastructure of TNTs using Scanning and Transmission EM (SEM and TEM, respectively) has only been analyzed in a handful of articles ^1,15-18^.

Although very similar under fluorescence microscopy (FM), TNT formation appears to be oppositely regulated by the same actin modifiers that act on filopodia ^19^. Furthermore, filopodia have not been shown to allow cargo transfer ^13,20,21^. Thus, we hypothesize that TNTs might be different organelles from filopodia and display structural differences in morphology and actin architecture.

In order to compare the ultrastructure and actin architecture of TNTs and filopodia at high resolution and ensure that the structures identified by TEM/SEM represent the functional units observed by FM, we employed a combination of live imaging, correlative light- and cryo-electron tomography (ET) approaches in two different neuronal cell models (mouse cathecholaminergic CAD cells and human neuroblastoma SH-SY5Y cells) ^19,22-25^.

We found that single bridging connections by FM are in most cases made up of a bundle of individual TNTs (iTNTs), each surrounded by a plasma membrane and containing its own highly organized parallel actin bundle. In addition, we also demonstrate that iTNTs contain vesicles, mitochondria, and other membranous compartments that appear to be traveling along actin filaments unidirectionally. Finally, by using correlative FIB-SEM we show that TNTs can be open on both ends, thus challenging the Dogma of a cell as an “individual unit” ^26^.

Collectively, our observations provide the first description of TNTs at the structural level and demonstrate that TNTs are different structures from other actin-containing cellular protrusions such as filopodia. We also describe an imaging work-flow that, although technically challenging, better preserves these fragile structures in native conditions, and is useful for studying their structure and the role they may play in other cell types and physiological models.

## Results

### Structural analysis of TNTs using cryo-correlated approaches

To study the ultrastructure of TNTs connecting CAD cells (**Supplementary Fig. 1A**) ^19,22^ we applied correlative light- and electron microscopy. Initial attempts of analyzing TNTs by correlative SEM showed that most TNTs connecting CAD cells broke during sample preparation (**Supplementary Fig. 1C–1D)** and only few, thick, more robust TNTs were preserved (**Supplementary Fig. 1E**) ^1,15,16^. Intriguingly some of the broken structures appeared to be comprised of multiple smaller tubes (yellow arrowheads, **Supplementary Fig. 1F**).

To better preserve these thin and fragile membranous structures, cells were labeled for FM, fixed by rapid freezing, and imaged at nanometer resolution by correlative cryo-FM and cryo-TEM/ET under fully hydrated conditions (see workflow detailed in **Fig. 1A)**. FM was used to screen for long (≥10 μm) and direct cell-to-cell connections labeled with wheat-germ agglutinin (WGA) hovering over the substrate, a phenotypic criteria used to identify TNTs ^1,2,19,23,27^.

**Figure 1.**
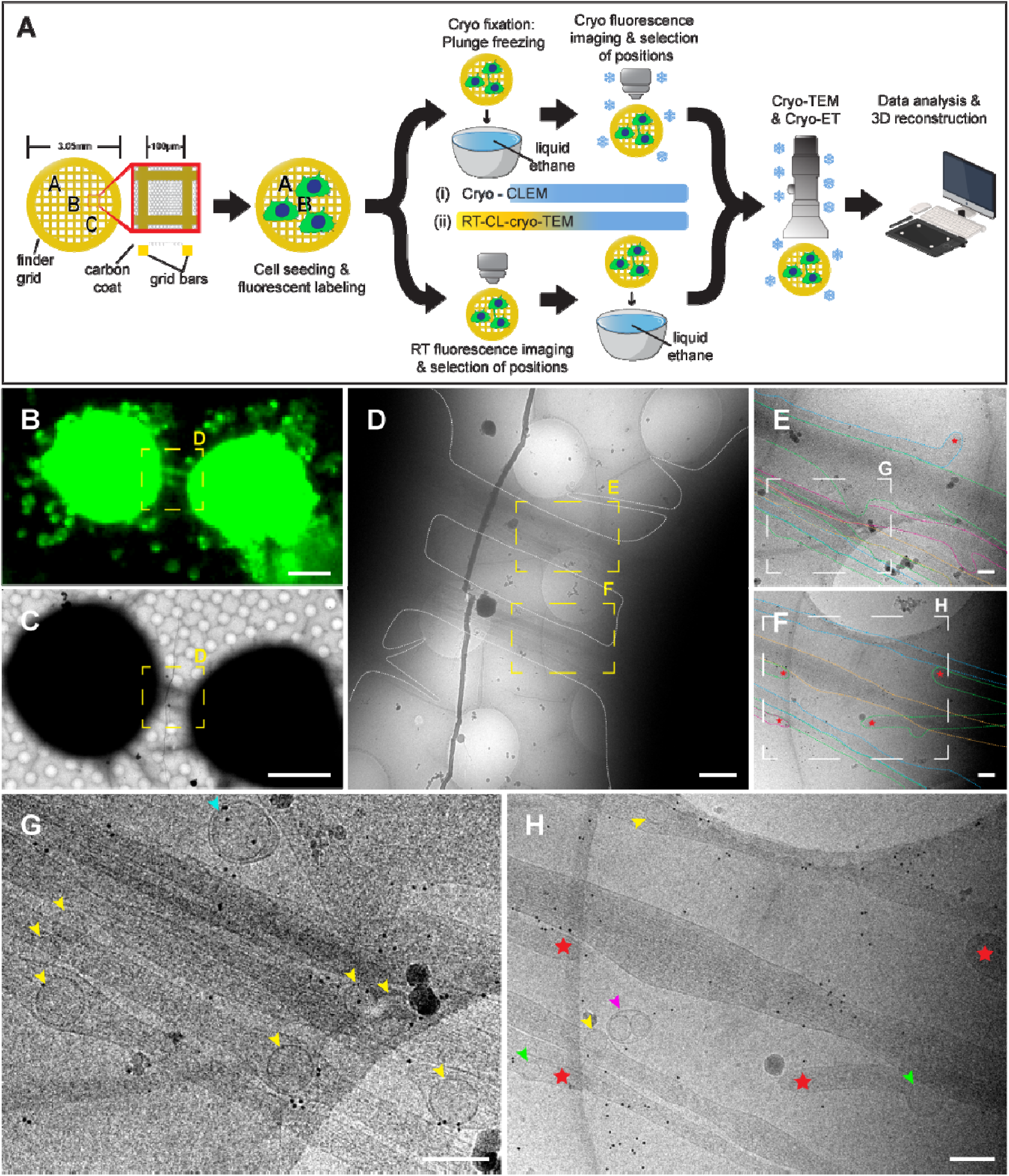
Structural Imaging of TNTs Using Correlated Light and Cryo-Electron Microscopy Strategies. (**A**) A schematic diagram of the experimental workflow and approaches used to observe TNT-connected CAD cells by cryo-TEM. (**B**) Cryo-FM image of two CAD cells connected by a TNT stained with WGA (green). (**C**) Cells in **B** observed in cryo-TEM at low magnification. Yellow dashed squares in **B**–**C** are shown at intermediate magnification in **D**. Yellow dashed rectangles in **D** are shown at high magnification in **E** and **F**, respectively. The plasma membrane of individual TNTs (iTNTs) were drawn with dotted colored lines in **E** and **F** to show membrane boundaries. Enlargements of white dashed rectangles in **E** and **F** are shown in **G** and **H**, respectively. (**G**–**H**) Various vesicular compartments were observed within and between iTNTs: yellow arrowheads show single vesicles; pink arrowheads show vesicles surrounded by an uter membrane; turquoise arrowheads show vesicles enclosed by a double membrane. Red stars in **E**, **F**, and **H** show tips of iTNTs extending/retracting from neighboring cells. Green stars indicate vesicles inside extending/retracting iTNT tips observed in **H**. Scale bars: **B**, **C**, 10µm; **D**, 1µm; **E–H**, 200nm.

Two correlative light and electron microscopy (CLEM) approaches were employed to image cells grown on grids: i-CLEM, and ii-CLEM. By i-CLEM, most TNTs that appeared as one tube under FM and transmitted light (**Fig. 1B–1C**) were comprised of a bundle of individual tunneling nanotubes (iTNTs), each delimited by its own plasma membrane as suggested by our SEM images (**Supplementary Fig. 1F**). iTNTs ran mostly parallel to and occasionally braided over each other (**Fig. 1D–1H**). We also observed that their tips could come from opposite directions, suggesting that they were in the process of extending to other cells or retracting from opposite cells (red stars, **Fig. 1F** and **1H**). Single thicker TNTs (600-900 nm in diameter), were rarely observed (**Supplementary Fig. 2A**).

TNTs have previously been reported to transfer vesicles, organelles, and various cargoes between cells by FM and live imaging ^2,4,5,22^; however, whether these cargoes were transported through the TNT lumen or by surfing on the limiting membrane could not be shown due to resolution limitations. By cryo-EM, we were able to observe vesicular membrane compartments located inside and between iTNTs. These vesicles were heterogeneous in size and shape, spanning from single membrane spherical vesicles (yellow arrowheads, **Fig. 1G–1H**, **2A–2E**, **2G**) with an average size of 109.12 nm (SD=25.55 nm) (**Fig. 2I**), to multi-vesicular compartments (blue arrowheads, **Fig. 2F** and **2H**). This observation supports our previous FM studies in CAD cells showing TNT-mediated transfer of DiD-labeled vesicles, lysosomes, and aggregated proteins between cells ^7,22,27^, and indicates that transfer occurs through the TNT lumen. Intriguingly, we also observed vesicles inside iTNTs with tips having opposite orientations (green arrowheads indicate vesicles, and red stars show the tips of tubes, **Fig. 1H**). This finding suggests that bidirectional cargo transfer observed by FM in one TNT could result from unidirectional transfer of vesicles inside iTNTs running in opposite directions. On the other hand, external vesicles were enclosed by a double membrane (turquoise arrowheads, **Fig. 1G**, **2E**, and **2H**), suggesting that they resulted from single-membrane vesicles budding off the plasma membrane of an iTNT. Several vesicles coupled together surrounded by an outer membrane were also observed outside iTNTs (pink arrowheads, **Fig. 1H** and **2B–2C**) which might also be explained by a membrane pinch-off or a break of the iTNT containing them.

**Figure 2.**
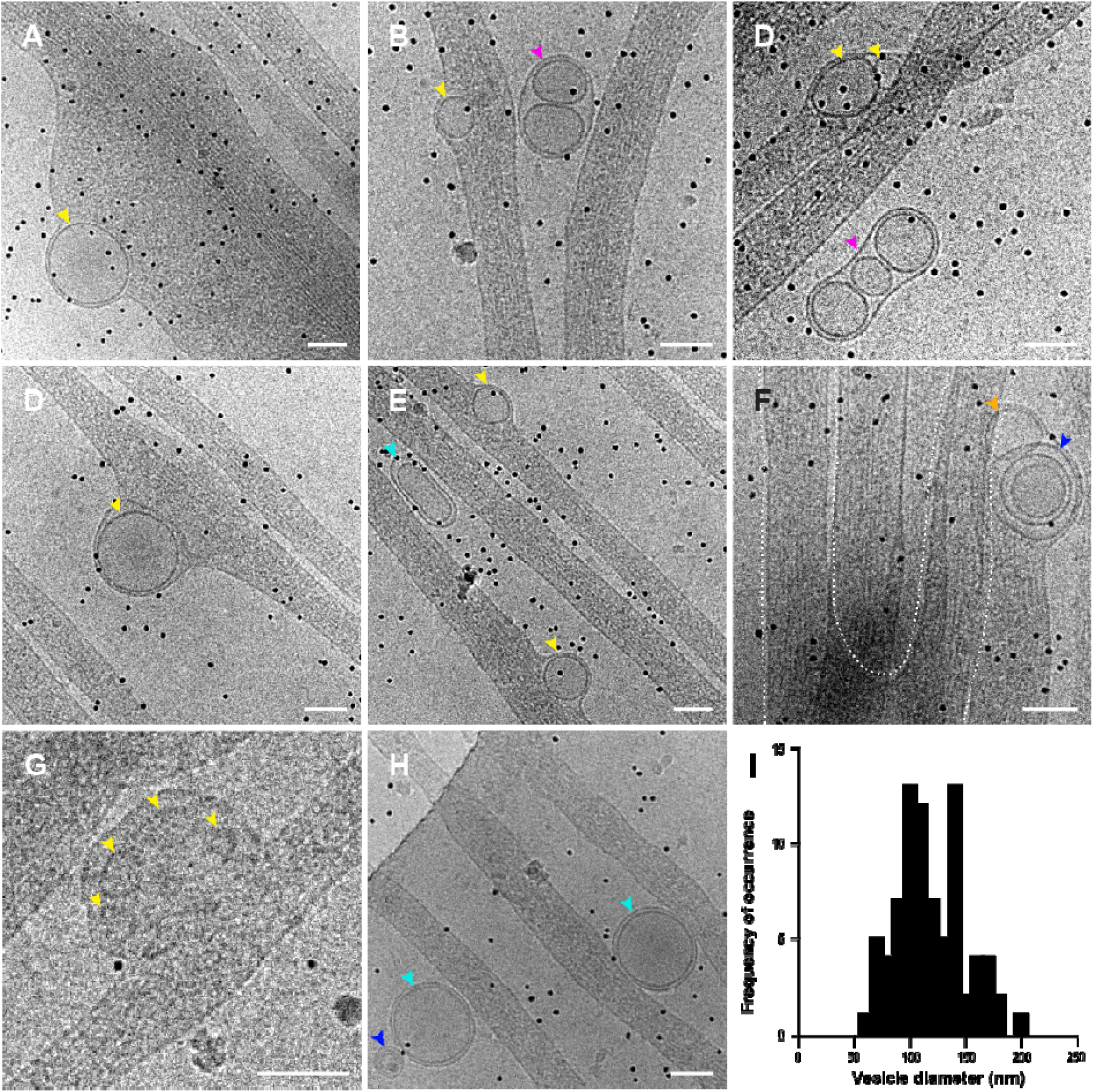
Gallery of Different Membrane Compartments Observed Outside and Within iTNTs. (**A–H**) Representative cryo-electron micrographs of membrane compartments and vesicular structures associated with iTNTs. Blue arrowheads in **F** and **H** indicate vesicles with a multi-lamellar appearance. Yellow arrowheads in **A**–**G** show single vesicular compartments. Turquoise arrowheads in **E**–**H** show extracellular vesicles with a double membrane. Pink arrowheads in **B**–**C** show groups of vesicles surrounded by a secondary membrane. Vesicles squeeze actin bundles together and create a bump on the plasma membrane (**A**–**G**). (**I**) Distribution of the diameter of vesicles in nm. Scale bars: **A–H**, 100nm.

Overall, by cryo-correlative light- and electron microscopy we discovered novel structures that correspond to the FM definition of TNTs. However, the number of TNTs imaged by cryo-CLEM was not sufficient for quantitative analysis, therefore we established a variant work flow by chemically fixing CAD cells and imaging them by FM at room temperature prior to rapid freezing and cryo-TEM (ii-CLEM) (**Fig. 1A**). Importantly, by cryo-EM, structural details looked indistinguishable between cryo- and chemically fixed samples (i.e. i-CLEM vs. ii-CLEM) (**Supplementary Fig. 3A–3B**) ^28^.

### Quantitative ultrastructural analysis of TNTs using cryo-electron tomography

To increase the number of TNTs on TEM grids even further, we turned to our previous findings showing that TNTs and filopodia are regulated by the same actin modifiers but in an opposite manner ^19^. CK-666, an Arp 2/3 complex inhibitor previously shown to inhibit filopodia ^29^, was tested for its effect on TNT formation. By using a method reported before ^19,30^ we quantified filopodia by counting vinculin-labeled peripheral spots in CAD cells. As hypothesized, our results show that CK-666 inhibits the formation of adherent filopodia (**Supplementary Fig. 4A–4B**), while the same treatment increased the number of cells connected via TNTs (**Supplementary Fig. 4C–4D**). These CK-666 induced TNTs were functional as assessed by a transfer assay previously used to monitor the transfer of DiD-labeled vesicles between a donor-acceptor cell co-culture ^19,23,27^, which shows a significant increase in the percentage of acceptor cells containing DiD-vesicles, (**Supplementary Fig. 4E–4F**). Importantly, TNTs in CK-666 treated CAD cells showed no detectable structural differences in cryo-TEM compared to the untreated control. (**Supplementary Fig. 4G–4H**).

We used these conditions to elucidate the spatial arrangement of iTNTs with respect to each other and to extract high-resolution detail of the actin organization and the vesicular compartments located inside iTNTs by cryo-ET. TNT-bundles had a diameter between 145 and 700 nm (305 nm on average) and were comprised of 2 to 11 iTNTs with an average diameter of 122.61 nm (SD= 65.73 nm). While the majority of iTNTs (~95%) measured up to 200 nm in diameter (**Fig. 3J**), thicker iTNTs (up to 550 nm) were occasionally observed. The spacing between individual tubes ranged between 8 and 90 nm for parallel iTNTs, while for less parallel iTNTs the spacing increased to 200 nm. 90% of these distances in parallel iTNTs ranged between 8 and 60 nm (**Supplementary Fig. 5**).

**Figure 3.**
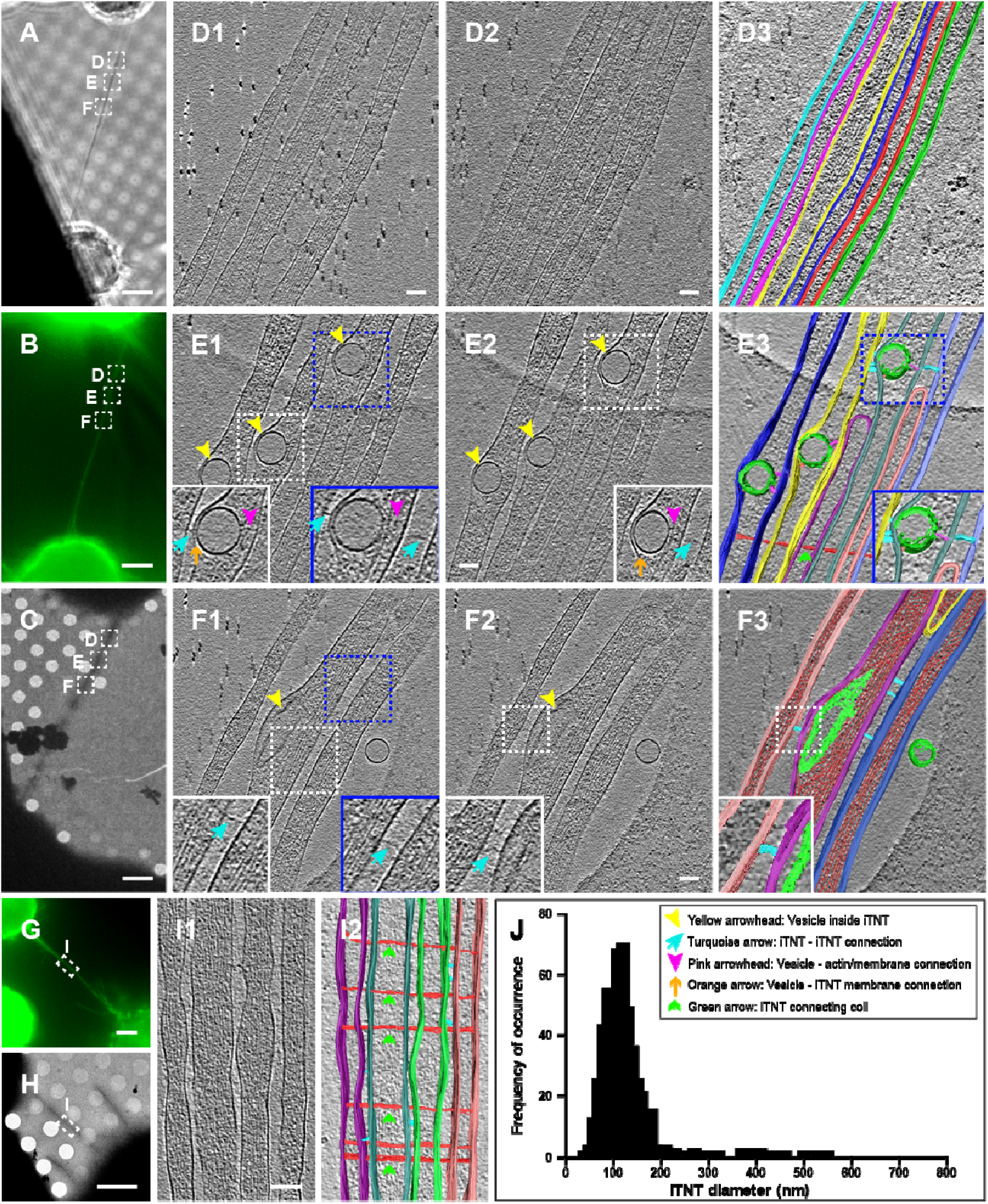
Imaging iTNTs in 3D Using Correlated Light and Cryo-Electron Tomography Approaches. CAD cells connected by a TNT stained with WGA (green) were imaged by phase contrast (A), epifluorescence (**B**), and low magnification TEM (**C**). Dashed squares over the TNT in A– **C** correspond to high magnification cryo-ET slices (**D1**–**D2**, **E1**–**E2**, **F1**–**F2**, and Supplementary Videos 1–3). (D3) Rendering of D1. (**E3**, **F3**) Renderings of tomograms (Supplementary Videos 2– 3). Turquoise arrows in **E1**–**E2** and **F1**–**F2** show filaments connecting iTNTs. Vesicles within iTNTs (yellow arrowheads, **E1**–**E2**, **F1**–**F2**) use thin filaments to connect to the plasma membrane on one side (orange arrows, **E1**–**E2**) and actin on the other (pink arrowheads, **E1**–**E2**). Additional example of cells connected by a TNT is shown in **G** (epifluorescence) and H (low magnification TEM). Region denoted by a white dashed rectangle in **G**–**H** is shown at high magnification cryo-TEM in **I1**. (**I2**) Rendering of I1 (Supplementary Video 4). A thin filament connects the bundle of iTNTs together by coiling around (green arrows, **E3**, **F3**, and I2). (J) Distribution of the diameter of iTNTs in nm. Scale bars: **A**–**C**, **G**–**H**, 5µm; **D**–**F**, **I**, 100nm.

By cryo-ET (**Fig. 3A–3C and 3G–3H**), iTNTs displayed thin structures connecting two iTNTs (turquoise arrows, **Fig. 3E**, **3F**, and **Supplementary Videos 2–4**) occasionally occurred along the entire length of iTNTs giving a ‘pearling’ effect (**Fig. 3I** and **Supplementary Videos 4–5**). Finally, our tomograms suggest that iTNTs were held together by long threads that coiled around them (green arrows, **Fig. 3E3** and **3I2**). While we could not investigate the nature of these formations, we also observed them in native frozen conditions, suggesting that they are not artifacts resulting from chemical fixation.

Another open question in the field is how cargo moves inside TNTs. We observed that vesicles inside iTNTs were typically located between the plasma membrane and a parallel actin bundle. Because their size exceeded the diameter of the individual tube, they bulged out from the plasma membrane, squeezing the actin filaments (**Fig. 1G–1H**, **2A–2F**, and **3E–3F**). By cryo-ET, we observed thin electron-dense structures that presumably connected intracellular vesicles to actin filaments (pink arrowheads, **Fig. 3E**), and short (~10 nm) spike-like structures that seemed to link vesicles to the plasma membrane on the opposite side of the actin bundle (orange arrows, **Fig. 3E**). These observations are consistent with the hypothesis that vesicles move inside TNTs on actin filaments using myosin motors ^1,27^.

Within iTNTs, actin was organized in long bundles of filaments running parallel to each other (**Fig. 3D–3F**, **3I**, **4A**, and **Supplementary Videos 1–5**). Due to beam damage, a technical drawback inherent to cryo-TEM, we were not able to image the actin bundle along the entire length of the TNT. However, by imaging non-consecutive (1.2-1.5 µm) regions of TNTs, each iTNT contained an uninterrupted parallel actin bundle extending along the whole length of the area imaged. This suggests that each iTNT contains a single continuous bundle of parallel actin filaments (**Fig. 3D–3F)**. Actin bundles filled the entire lumen of with a diameter of < 300 nm iTNTs (**Fig. 1G–1H**, **2A–2H**, **3**, and **4A–4B**). Tracing plot profiles and performing the Fourier power spectra analysis on our cryo-CL-ET images estimated the average distance between the centers of actin filaments in bundles at 10.05 nm (SD= 0.84 nm) with an inter-filament distance of 4.7 nm (SD=1.08 nm) (**Fig. 4A–4F**).

**Figure 4.**
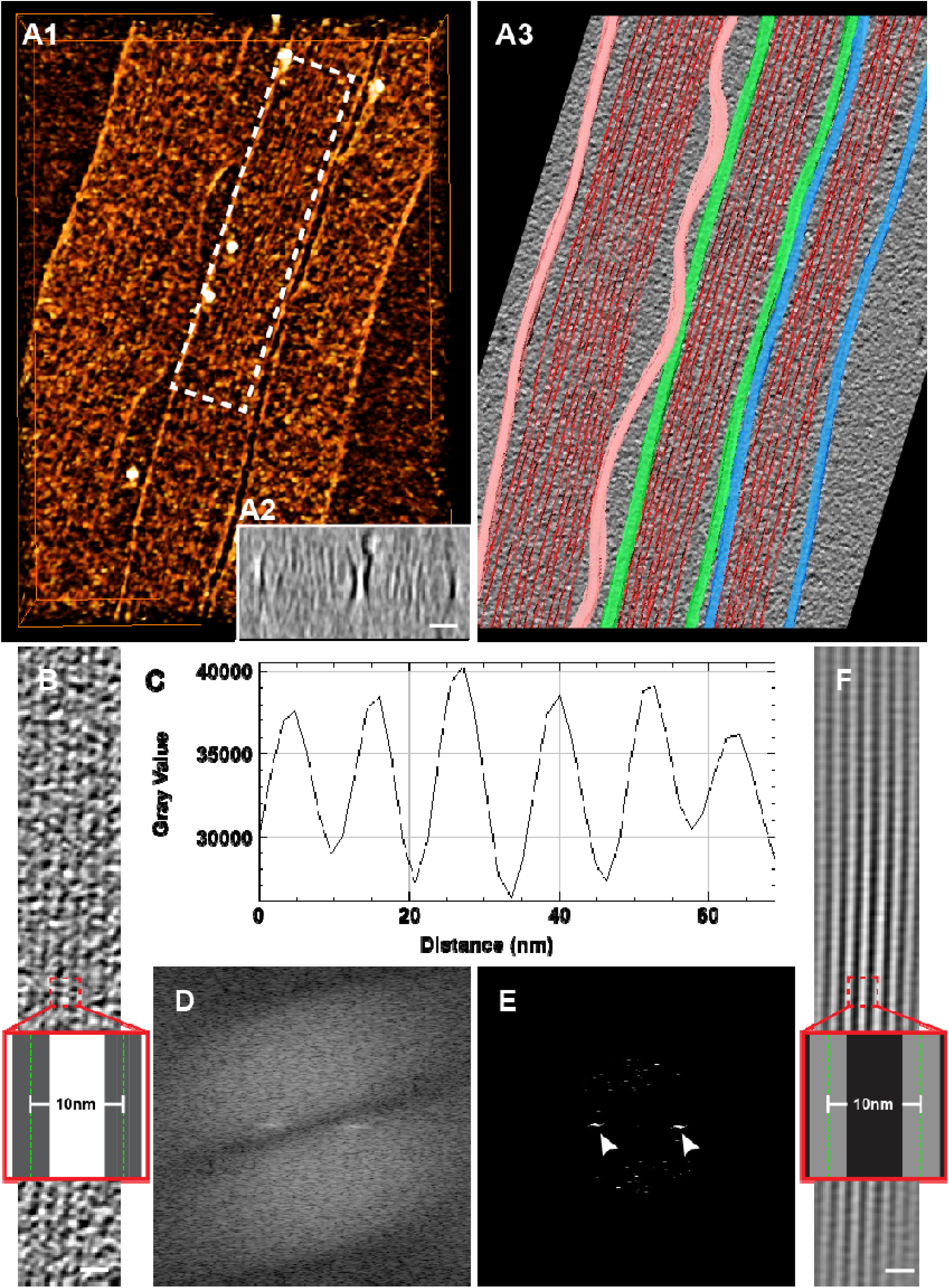
Cryo-Electron Tomography and Image Analysis Reveal F-actin Organization in iTNTs. (**A1**) Surface rendering obtained from a tomogram (Supplementary Video 5) showing F-actin bundles within iTNTs. (**A2**) Cross-section through the cryo-electron tomogram in **A1** showing two iTNTs. (**A3**) Rendering of tomogram used to obtain **A1**. (**B**) Tomogram slice corresponding to white dashed rectangle in **A1**. The average size and spacing between filaments of an iTNT shown in **B** were measured by computing the plot profile of actin bundles residing within. (**C**–**E**) Filament-to-filament distance measured by extracting peaks in the frequency domain using the fast Fourier transform (white arrowheads, **E**) and by computing plot profiles (**C**). (**F**) Model of the actin filaments displaying a parallel bundle in iTNTs obtained by retransforming peaks in the frequency domain shown in **E**. Insets in **B** and **F** show distances between the average measurements in nm obtained by the analysis in **C** and **D**. Scale bars: **A2**, **B**, **F**, 20nm.

### Comparing and contrasting TNTs to filopodia

The structures we described above as TNTs appeared different compared to filopodia previously studied by cryo-ET ^31,32^. Thus, we decided to use our cryo-EM approach to analyze the structure of filopodia in CAD cells in order to directly compare them with TNTs in the same cell type (**Fig. 5A–5F**). To increase filopodia number on TEM grids, CAD cells were transfected with the vasodilator-stimulated phosphoprotein (VASP), a protein previously shown to be an effective inducer of filopodia in different cells ^13,19,27^. Filopodia induced by VASP looked indistinguishable from those of untreated cells at the ultrastructural level (**Fig. 5A–5F**).

**Figure 5.**
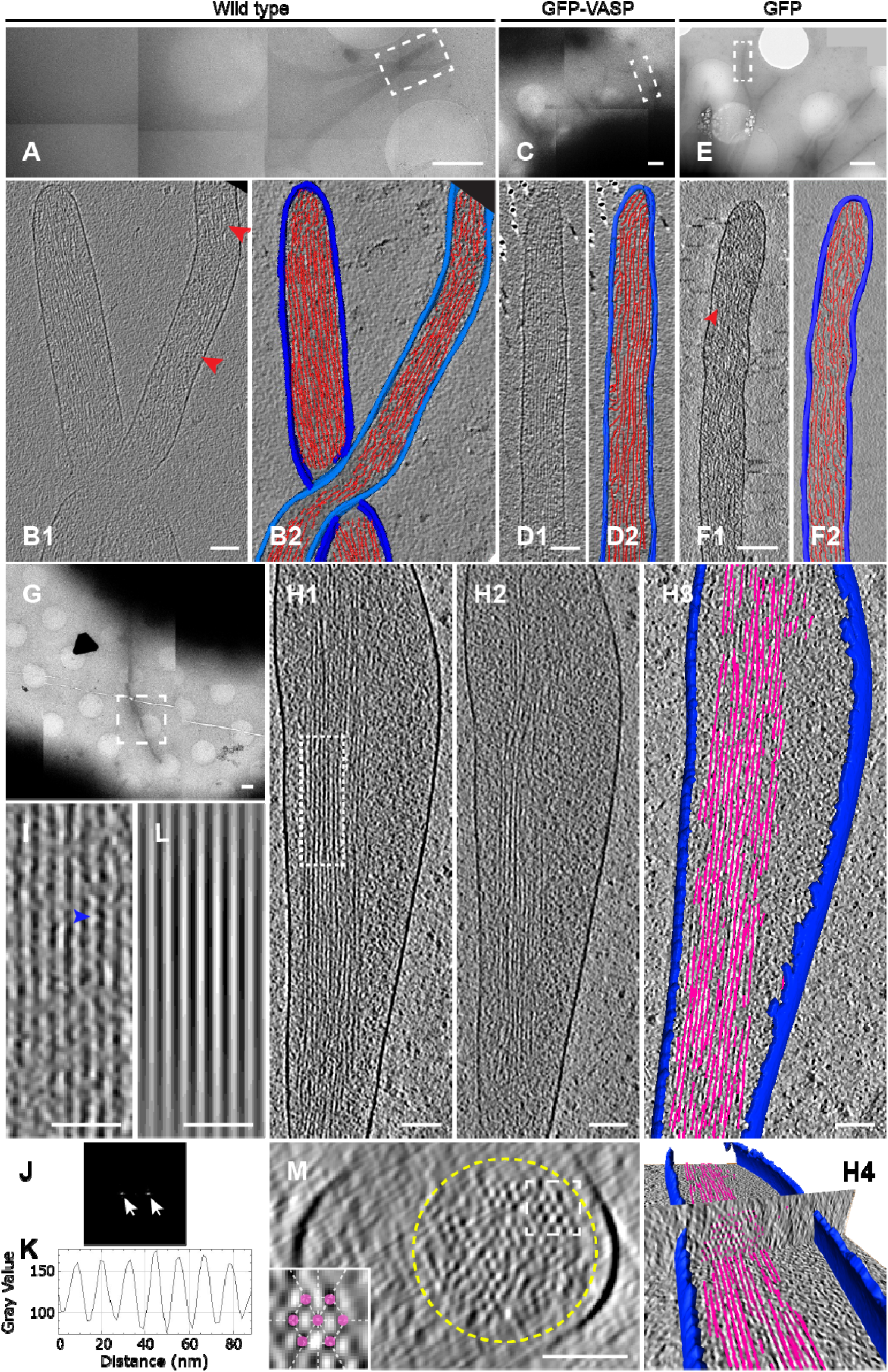
Cryo-Electron Tomography and Image Analysis Reveal Actin Ultra-Structure in Filopodia. Representative electron micrographs of filopodial protrusions of untreated (**A**–**B**, **G**–**H**), GFP-VASP (**C**–**D**), and GFP (control) (**E**–**F**) transfected CAD cells imaged by cryo-TEM at low magnification **(A, C, E, G)**. (**B1**, **D1**, **F1**) Slices from volumes rendered in **B2**, **D2**, and **F2** correspond to white dashed rectangles shown in **A**, **C**, and **E**, respectively. Red arrowheads in **B1** and **F1** indicate branched actin configuration. (**H1**–**H2)** slices corresponding to white dashed square in **G**. (**H3**–**H4**) 3D automated rendering of **H1**–**H2** and cross-section of the filopodium (**H4**). Segmentation of the plasma membrane (blue) and actin (pink). (**I**) High magnification view of filaments corresponding to the white dashed rectangle in **H1**, where filaments display a wavy periodic pattern (blue arrowhead, **I**). By computing its plot profile (**K**), the average filament size and spacing between filaments was measured. (**L**) Model of the parallel filament bundle obtained from the peak extraction in the fast Fourier transform (**J**). (**M**) Cross-section through the cryo-electron tomogram of the white dashed rectangle in **G.** Scale bars: **A**, **C**, **E**, 1µm; **B**, **D**, **F**, **H**, **M**, 100nm, **G**, 1µm; **I**, **L**, 50nm.

Filopodial extensions in CAD cells exhibited lengths that span between one and several microns with an average diameter of 174.86 nm, (ranging between 118.29 nm and 285.5 nm). These measurements were close to those we obtained for TNTs, making the discrimination between these two structures challenging by FM. However, unlike TNTs, filopodia were single isolated protrusions that displayed both short (**Fig. 5B**, **5D**, and **Supplementary Videos 6–7**) and long conformations (**Fig. 5H** and **Supplementary Video 9**) that can be straight, bent, or twisted around other filopodia (**Fig. 5B** and **Supplementary Video 6**).

Actin filaments within the filopodia of CAD cells showed two different organizations. As previously described for other neuronal cell lines ^32^, actin filaments were mostly organized in tight parallel bundles that extended into the tip of the filopodium (**Fig. 5B**, **5D**, **5H**, **5I**, and **Supplementary Videos 6–7,** and **9**). Differently form TNTs, filopodia actin filaments did not run uninterrupted along the whole length of the area imaged (1.2-1.5 µm). Instead, they were organized in bundles comprised of shorter filaments with a length varying between 300 and 1100 nm, with only 15% having a length larger than 1 µm (**Fig. 5H**). Alternatively, filopodia also displayed short parallel actin bundles (**Fig. 5B**, **5H**, and **Supplementary Videos 6** and **9**) that intermingled with short-branched filaments (red arrowheads, **Fig. 5B**, **5F1,** and **Supplementary Videos 6** and **8**), never observed in TNTs, but similar to what was previously shown for filopodial conformations in Dyctostelium discoideum amoebae ^31^.

In agreement with results reported by Aramaki et al., in a different neuronal cell line ^32^, the average distance between actin filaments in bundles was 10.36 nm (SD=0.4 nm) with an inter-filament distance of 4.68 nm (SD=0.73 nm) (**Fig. 5I–5K**). This is similar to the estimate obtained for actin bundles within iTNTs (**Fig. 4C–4E**), suggesting that at the resolution level we achieved for TNTs, the actin arrangement in both structures was similar. We further analyzed cross-sections of filopodia and show that the actin bundle diameter ranged between 80 and 170 nm and the number of filaments per bundle ranged between 38 and 150. Moreover, actin filaments in filopodia were bundled by cross-linkers arranged in hexagonal arrays comprised of 12-30 filaments, with a ~4.7 nm spacing (**Fig. 5M** and **5H4**). A similar actin-bundling hexagonal pattern was previously shown to be mediated by the actin-binding protein fascin ^32^.

Finally, another difference with TNTs is filopodia’s general lack of intracellular vesicular structures (as previously reported in other cell types) ^31,32^.

### SH-SY5Y cells connect via TNTs capable of transporting mitochondria

To confirm that the structural characteristics of TNTs are not cell-type dependent, a neuronal cell model of human origin, SH-SY5Y cells, previously shown to connect via TNTs using FM, was analyzed next ^4,24,25^ (**Supplementary Fig. 1B**). To investigate the nature of TNTs connecting SH-SY5Y cells we first analyzed them by SEM. Similar to CAD cells, thicker TNTs appeared to endure harsh classic EM sample preparation steps (red arrowhead, **Supplementary Fig. 1G**), while thinner structures would often break (yellow arrowheads, **Supplementary Fig. 1G** and **1H**). Compared to CAD cells, however, more thin structures connecting SH-SY5Y cells survived the classical embedding conditions (blue arrowheads, **Supplementary Fig. 1I**). By employing CLEM approaches on this cell model, TNTs connecting SH-SY5Y were also comprised of bundles of two or more iTNTs (**Fig. 6A–6D**) and only occasionally single thick connections were observed (**Supplementary Fig. 2B**). iTNTs connecting SH-SY5Y cells had an average diameter of 120.71 nm (SD= 71.39 nm) (**Fig. 6E)** and contained vesicular compartments (yellow arrowheads, **Fig. 6C–6D**), with an average diameter of 104.04 nm (SD=57.78 nm) (**Fig. 6F**). Importantly, control experiments demonstrated that iTNTs connecting SH-SY5Y cells fixed by rapid freezing were virtually identical to those chemically fixed, as in CAD cells (**Supplementary Fig. 3C–3D**).

**Figure 6.**
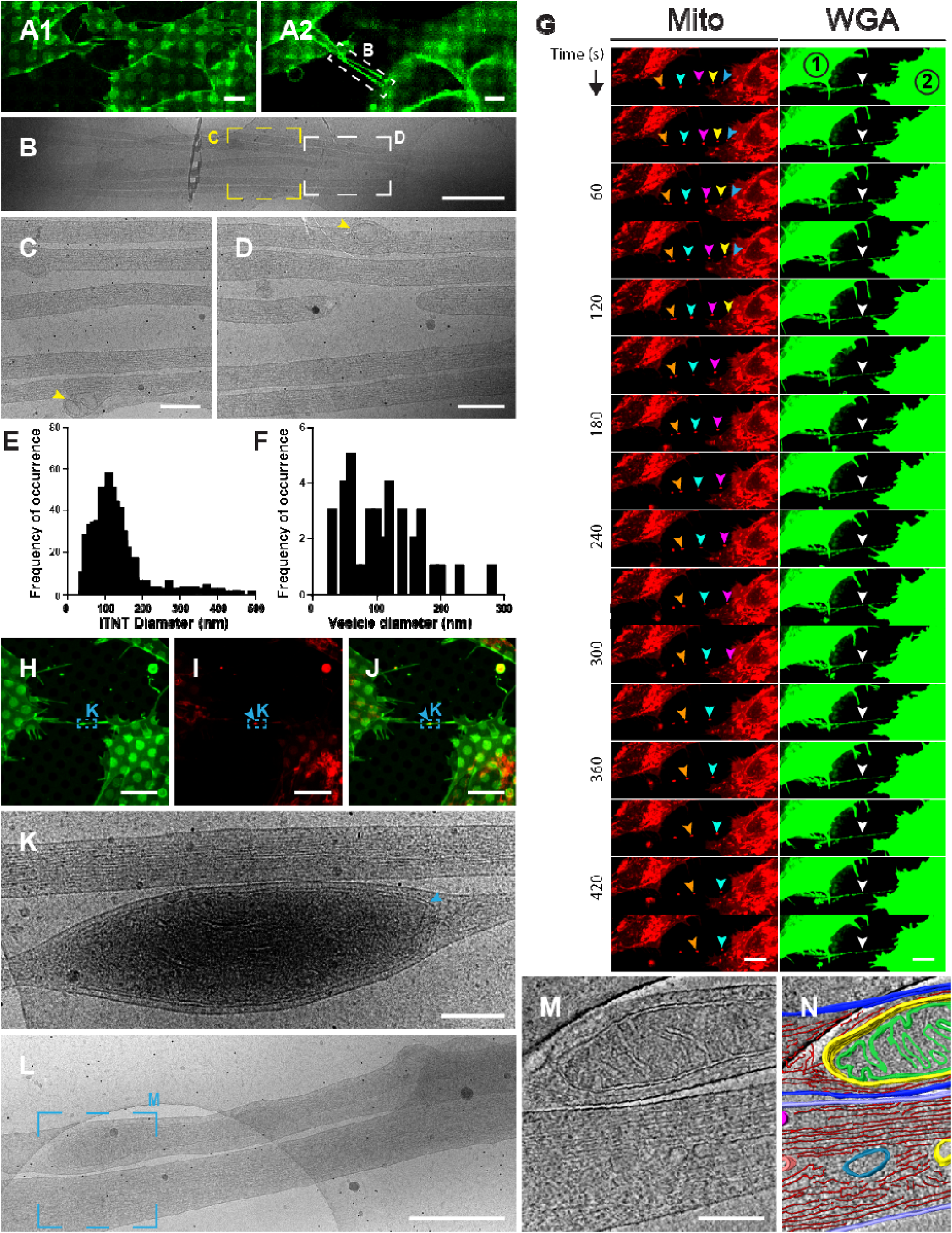
Structural Analysis and Live Imaging Reveal Mitochondrial Transport via iTNTs. (**A**) Confocal micrograph of SH-SY5Y cells stained with WGA (green). (**A1**) Lower and (**A2**) upper optical slices show TNTs hovering over the substrate. (**B**) Low-magnification electron micrograph corresponding to white dashed rectangle in **A2**. (**C**–**D**) High-magnification micrographs corresponding to yellow and white dashed squares in **B**. Yellow arrowheads indicate membranous compartments inside iTNTs. (**E**–**F**) Diameter distribution of iTNTs (**E**) and vesicles found within them (**F**) in nm. (**G**) Time-lapse sequence of SH-SY5Y cells stained with WGA (green) and MitoTracker (red) reveal mitochondrial puncta (orange, cyan, pink, yellow, and blue arrowheads) traveling across a TNT (white arrowhead) entering the apposing cell, (cell #2). (**H**–**J**) SH-SY5Y cells stained with WGA (green) and mitotracker (red), (blue arrowheads indicate mitochondria inside the TNT). (**K**) Micrograph corresponding to blue dashed rectangles in **H**–**J** reveal mitochondria (blue arrowhead) observed by fluorescence (**H**–**J)**. (**L**) Micrograph displays two iTNTs, one containing mitochondria. (**M**) Tomogram slice corresponding to **L** (blue dashed square) reveal mitochondrial cristae, actin, and membranous compartments found inside iTNTs. (**N**) Rendering of **M**. Scale bars: **A**–**B**, 5µm; **C**, 1µm; **D**, **K**, **M**, 200nm, **G**–**J**, 10µm; **L**, 500nm.

In order to demonstrate that the structures identified by cryo-CLEM in SH-SY5Y cells were functional we analyzed whether they also transferred cargo. In addition to vesicles, TNTs are able to transfer mitochondria between connected cells ^4,33-35^. Compared to DiD-vesicles, mitochondria are particularly bright when fluorescently labeled and therefore well suited to be imaged by live microscopy, ^36,37^. When cells labeled with WGA (green) and MitoTracker (red) were recorded by live cell imaging mitoTracker-positive puncta moved inside a TNT in a unidirectional fashion at an average velocity of 0.0514 µm/sec (SD=0.0261 µm/sec) (**Fig. 6G** and **Supplementary Video 11**). As previously reported for PC12 cells ^36^, mitochondria did not move through the TNT at a uniform speed and seemed to accelerate in segments where the TNT appeared to be completely straight. As exemplified in **Supplementary Video 12**, labeled mitochondria traveled all along an iTNT and accumulated in the cytoplasm of the neighboring cell (blue arrowhead, **Supplementary Fig. 6A** and **Supplementary Video 12).**

The presence of mitochondria inside iTNTs was confirmed at the ultra-structural level by ii-CLEM and cryo-ET. By cryo-ET, the mitotracker labeled structures seen by FM (**Fig. 6H–6J)** were mitochondria based on their size, shape, and presence of mitochondrial cristae (**Fig. 6L–6N)**. Intriguingly mitochondria were observed in only one of the iTNTs which contained actin and often appear to bulge with the passage of mitochondria (**Fig. 6K**, **M**, **N**, and **Supplementary Fig. 6B–6I)**. Altogether, this data demonstrates that the structure of functional TNTs (i.e. allowing transfer of cargoes inside their lumen) is similar between CAD and SH-SY5Y cells, two different neuronal cell models from different origins and species.

### SH-SY5Y cells connect via continuous and closed-invaginating connections

Given that vesicles and larger organelles such as lysosomes and mitochondria are transferred between cells connected by TNTs, it is conceivable that TNTs are structures open on both ends connecting the cytoplasm of two neighboring cells. This notion, however, is still controversial as it has not been sufficiently supported by prior ultra structural data ^1,12,15^. Cryo-ET, was not suitable to address this question as connections between TNTs and cell bodies, were too thick (> 500nm in thickness) to be investigated by TEM (**Supplementary Fig. 7**). Therefore, we used focus ion beam SEM (FIB-SEM) tomography to analyze specifically the ‘contact zones’ where TNTs contacted the cell bodies of connected cells.

Contact zones were identified by FM (**Fig. 7A** and **7E**) and cells subsequently prepared for FIB-SEM tomography. Under the imaging conditions used, the resolution was sufficient to discern plasma membrane boundaries of cells and TNTs, which were segmented to visualize contact zones in 3D. The 3D reconstruction illustrated in **Fig. 7D** revealed two single contiguous TNTs with an average diameter of 176.3 nm (SD=18.65 nm) and 4.3 μm of length) crossing each other along the way and connecting a pair of cells (**Fig. 7C–7D**, **Supplementary Video 14**). In a separate example, we observed a pair of tubes with 220 nm in diameter (SD=35.72 nm) connecting two cells by FM (**Fig. 7E**). One of these tubes is inserted inside the apposing cell and appears to be closed at its tip (red arrowhead, **Fig. 7E–7F, Supplementary Video 15**). This invaginating event could presumably be the result of a pre- or post- TNT (fusion) event. Alternatively, this observation could also indicate that TNTs can either be open- or close- ended at contact zones. As illustrated in **Fig. 7F**, the imaging planes from FIB-SEM (**Fig. 7G**) correspond to fluorescent counterparts. It should be noted that TNTs in our FIB-SEM data did not display a straight and smooth shape typically observed by FM and cryo-EM. We believe that tubes can deform during sample preparation or milling. Nonetheless, to our knowledge this represents the first demonstration that TNTs observed by FM can be open-ended on both sides, thus directly linking the cytoplasm of two connected cells.

**Figure 7.**
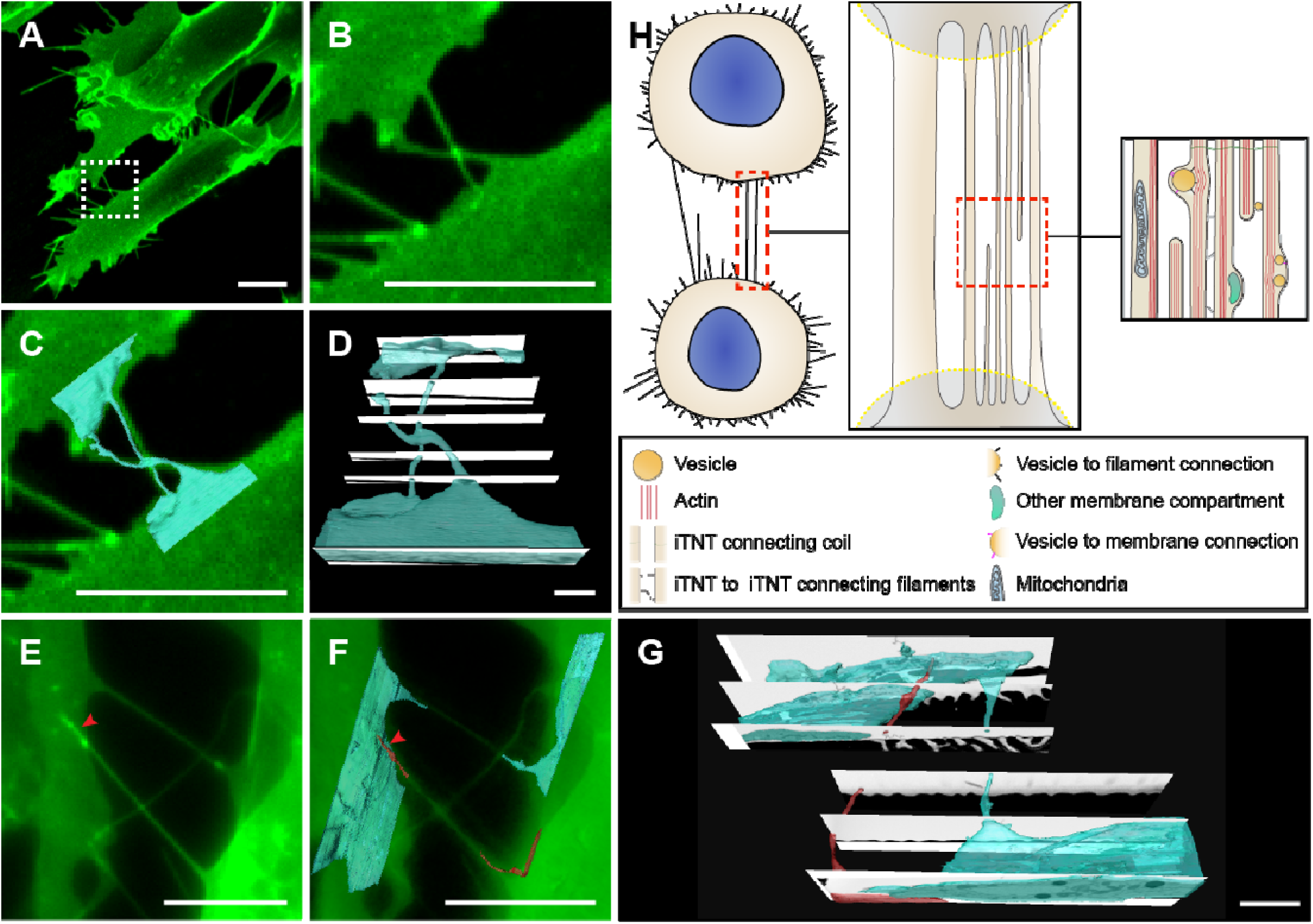
FIB-SEM of TNTs in SH-SY5Y Cells Reveals Open-Ended Contact Zones. (**A, E**) Confocal micrograph of SH-SY5Y cells stained with WGA (green). (**B**) Magnified view of white dashed box in **A**. (**C**) Overlay of 3D rendering of FIB-SEM tomogram segmentations over fluorescent counterpart, **A,** reveal open-ended TNTs connecting SH-SY5Y cells. (**D**) 3D rendering of FIB-SEM tomogram segmentation shown in **C**. (**F**) Overlay of 3D rendering of FIB-SEM tomogram segmentations over fluorescent counterpart, **E,** reveal two connections: one open-ended TNT, and one invaginating inside the cytoplasm of the apposing cell (red arrowhead). (**G**) 3D rendering of FIB-SEM tomogram segmentation shown in **F**. (**H**) Schematic diagram depicting how two cells connect via TNTs. TNTs can either be single thick connections or a bundle of thin individual TNTs (iTNTs). iTNTs contain vesicles and mitochondria. Membranous compartments within iTNTs use thin filaments to connect to actin on one side and to the inner side of the plasma membrane on the other. Thin membrane threads coils exist between and around several iTNTs. Presumable appearance of TNT ends, or contact zones, are indicated by light gray semicircles with a dashed yellow outline. Scale bars: **A**–**C**, **E**–**F**, 10µm; **D**, 2µm; **G**, 3µm.

## Discussion

Tunneling Nanotubes (TNTs) have been shown to play a vital role in the spreading of different cargos, organelles, and amyloidogenic proteins between cells *in vitro* and have therefore been implicated in many different physiological and pathological processes ^2-5,22^.

While the morphological features of TNTs have been extensively described in various cell types using fluorescence microscopy, to date, no reliable EM method has been reported that would allow for the observation of TNTs at the nanoscale resolution without jeopardizing their fragile membranous and cytoskeletal composition. Indeed TNTs have only been studied at the ultrastructural level by conventional SEM and TEM methods in a handful of publications ^1,15-18^.

Here we set up a workflow for correlative light- and cryo-ET microscopy that, to the best of our knowledge, is the only approach to identify and characterize the precise structure of these cell-to-cell connections observed by FM. By using this approach, we were able to observe TNTs connecting two different neuronal cell models, CAD and SH-SY5Y cells at the nanometer resolution.

Our results show that most TNTs, which appear as single connections by fluorescence microscopy, are in fact made up of several individual tunneling nanotubes (iTNTs). iTNTs often run together in a parallel fashion but occasionally braid over each other. We speculate that this conformation might greatly increase their stability and elasticity, allowing them to withstand movements between connected cells. iTNTs contain actin bundles organized in a highly-ordered fashion and likely extend all along the tube. In the majority of cases, actin bundles filled the entire lumen within iTNTs suggesting the same actin polarity in a single tube. We also observed tips of iTNTs “likely” in the process of “extending to- or retracting from either of the two connected cells (**Fig. 1** and **6**). We hypothesize that iTNTs extending from opposing cells would have opposite polarities, thus allowing unidirectional transfer in opposite directions (always from donor to acceptor cell). In support of this hypothesis we could observe vesicles inside distinct iTNTs (within the TNT bundle) that appear to originate from opposite sides (**Fig. 1H** and **7H**). This could explain bidirectional vesicular transfer previously observed inside what appeared as a single TNT by FM. Further studies combining live microscopy with cryo-TEM will be required to directly confirm this hypothesis. Alternatively, one could imagine that each iTNT could contain actin filaments with opposite polarity, or both actin and microtubules. While our tomograms are suggestive of a single actin bundle per iTNT, the presence of tubulin has been observed (by FM) in thicker connections, mainly in cells of the immune system, but not inside TNTs from neuronal cells ^1,38,39^.

In addition, our tomograms suggest the existence of thin filaments that connect iTNTs (**Fig. 3** and **7H**). These contacts presumably increase the mechanical stability of the overall bundle and hold iTNTs together. Understanding whether these filaments are protein or lipid based will be important to decipher the molecular mechanism underlying the formation of iTNTs.

We also detected vesicular compartments of different shapes and sizes inside iTNTs (**Fig. 2**), reinforcing the wealth of literature demonstrating the intercellular transfer of vesicles via TNTs ^2,4,5^. In addition, we observed extracellular vesicles surrounded by a double membrane, which could have derived from either budding or fission from an iTNT. Future studies will be necessary to investigate both the nature of intra- and extracellular vesicles.

Nonetheless, by correlative cryo-TEM we identified mitochondria inside iTNTs connecting SH-SY5Y cells (**Fig. 6**). Together with live microscopy data showing mitochondria traveling through TNTs, these findings indicate for, the first time, that mitochondria can be transported within iTNTs in a tubulin-independent mechanism (**Fig. 6M**). Interestingly, while microtubules and their motors have been established as important factors for mitochondrial transport ^40^, emerging evidence indicates that mitochondria interact with the actin cytoskeleton in many cell types ^41-43^. Although we could track single mitochondria and measure their velocity by live imaging, the motors involved in their transport through iTNTs, as well as the motors involved in the transport of the different vesicular structures, remain to be identified.

Overall these data indicate that TNT are very different from filopodia. In order to directly demonstrate that they are different structures we applied our cryo-ET analysis and compared the two structures in CAD cells. Tomograms revealed that similar to other cells, filopodia in CAD cells are isolated protrusions with an average diameter of 200 nm and lengths between one and several microns. Filopodia also displayed various actin arrangements compared to TNTs, (either tight parallel bundles, but of shorter length compared to TNTs, or parallel bundles intermingled with short-branched filaments). Finally, as previously shown in other cell types ^31,32^, they do not appear to contain vesicles. Cross-section analysis in filopodia also showed a typical actin-bundling hexagonal pattern possibly due to the actin-binding protein fascin ^32^. Under the imaging conditions used, the number of filaments per bundle and actin bundling proteins could not be clearly detected for TNTs. This is possibly due to the lack of resolution achievable by our cryo-TEM on thick samples and the considerable technical challenges we faced when imaging iTNT bundles under cryogenic conditions compared to isolated filopodia. Thus, to further analyze the structural differences and similarities in the actin arrangements within iTNTs, a more powerful cryo EM approach would be needed (e.g. Titan/Krios TEM).

Next, to specifically address the controversial question in the TNT field regarding whether TNTs are connecting the cytosol of two cells, we performed FIB-SEM tomography, which allowed us to image the ends (or contact sites) of iTNTs. Interestingly, by FIB-SEM we were able to identify both open- and close-ended connections such as invaginations at contact zones. At this point, we cannot discriminate whether the differences observed are the result of the existence of distinct iTNTs or whether it is due to temporal pre or post fusion events. Nonetheless these data represent the first demonstration that open-ended TNTs exist and could correspond to the functional TNTs structures observed by FM.

Overall, the data presented here represent the first structural characterization of functional TNTs previously observed by light microscopy in two different neuronal cell models. They demonstrate their specific identity, highlight the morphological differences with filopodia, show the actin architecture at high resolution and the presence of vesicular structures and mitochondria inside them.

While many more questions remain open these data also guide the establishment of a TNT inducing and imaging platform for investigating the structural mechanism underlying TNTs, as well as for revealing the specific role that membrane and cytoskeleton-associated proteins may play in the formation and function of this important biological process.

## Materials and methods

### Cell preparation and transfections

CAD cells (mouse catecholaminergic neuronal cell line, Cath.a-differentiated) were cultured at 37°C in Gibco Opti-MEM (Invitrogen), plus 10% fetal bovine serum and 1% penicillin/streptomycin ^23^. SH-SY5Y cells (neuroblasts from human neural tissue) were cultured at 37 °C in RPMI-1640 (Euroclone), plus 10% fetal bovine serum and 1% penicillin/streptomycin. Transient transfections were performed with Lipofectamine 2000 (Invitrogen) in accordance with the manufacturer’s instructions. GFP-tagged constructs were used as previously described ^19^.

### Pharmacological treatments

#### Effect of CK-666 in the Number of Cells connected via TNTs and Vinculin Spots

For pharmacological assays, confluent CAD cells were mechanically detached and counted; 100,000 cells were plated for 24 hrs on Ibidi µ-dishes (Biovalley, France). Cells were then treated with CK-666 (SML0006, Sigma), an Arp 2/3 complex inhibitor, which was made up as 75 mM stock solution in DMSO and diluted in complete medium to a final concentration of 25 µM or 50 µM. DMSO-alone control treatments were performed as part of every experiment involving drug treatment. Cells were treated for 15 min at 37 °C before fixation and fluorescent labeling.

#### Effect of CK-666 in the Transfer of DiD-labeled Vesicles in Co-Culture System

Acceptor cells were plated in T25 flasks (Corning, USA) and transiently transfected with an H2B-mcherry vector after 24 hrs. The following day, donor cells were detached and resuspended in complete medium with 1 µM of DiD-labeled vesicles and incubated for 30 min at 37 °C. Donor cells were then washed with complete culture medium, mixed with acceptor cells (1:1), and plated on 35 mm Ibidi µ-Dish (Biovalley, France). After an incubation of 1h at 37 °C, cells were treated with CK-666 (25 µM and 50 µM) and DMSO (control) for 6hours. After washing cells with fresh, complete medium they were incubated at 37 °C O/N. On the next day, cells were washed with PBS, fixed in 2% PFA + 0.05% GA in 0.2 M Hepes for 15 min followed by fixation in 4% PFA in 0.2 M Hepes for 15 min, stained with DAPI (1:1000), and mounted with Aqua-Poly/Mount (Polysciences, Inc.). Images were acquired on an LSM 700 confocal microscope (Zeiss) with a 40X objective. The detection of DiD-labeled vesicles within the acceptor cells was analyzed as described in About S. et al., 2016 ^7^.

### Immunofluorescence labeling

CAD cells plated overnight were washed carefully, fixed with 4% paraformaldehyde (PFA) for 15 min at 37 °C, quenched with 50 mM NH4Cl for 15 min, permeabilized and blocked with 0.075% Saponin in PBS containing 2% BSA (w/v) for 1 hour at 37 °C. Cells were then incubated with a mouse anti-vinculin primary antibody (V9264, Sigma) at 1:500 in 0.01% saponin and 2% BSA (w/v) in PBS. Cells were thoroughly washed and incubated for 40 min with a goat anti-mouse AlexaFluor 488-conjugated secondary antibody (Invitrogen) at 1:500 in 0.01% saponin and 2% BSA (w/v) in PBS. Cells were then carefully washed in PBS and labeled with WGA-647 (1:300 in PBS) for 5 min at RT, and 5 min with Phalloidin-Rhodamine (1:300 in PBS); nuclei were labeled for 5 min with DAPI (1:1000). Cells were washed and mounted with Aqua-Poly mount (Polysciences, Inc.).

### Cell preparation for cryo-EM

Carbon-coated gold TEM grids (Quantifoil NH2A R2/2) were glow-discharged at 2 mA and 1.5-1.8×10^−1^ mbar for 1 min. Grids were sterilized under UV for 15 min (3X) at RT and incubated at 37 °C in complete culture medium. Grids were carefully washed in PBS (2X) and coated with 40 µg/mL of fibronectin for 20 min at 37 °C. After three PBS washes, 250,000 CAD and 700,000 SH-SY5Y cells were seeded on grids and incubated O/N at 37 °C, resulting in 3 to 4 cells per grid square. Prior to chemical or cryo-fixation, cells were labeled with WGA-Alexa-488 (1:300 in PBS) for 5 min at 37°C. For correlative light- and cryo-electron microscopy, cells were chemically fixed in 2% PFA + 0.05% GA in 0.2 M Hepes for 30 min followed by fixation in 4% PFA in 0.2 M Hepes for 30 min and kept hydrated in PBS buffer prior to vitrification.

*Cell vitrification.* Fluorescently labeled CAD and SH-SY5Y cells were rapidly frozen in liquid ethane ^44^ using a Leica EMGP system.

### Microscopy

#### Confocal Microscopy and Image Analysis

Fluorescent images from pharmacological assays were acquired using an inverted Zeiss LSM 700 confocal microscope. Quantification of TNT-connected cells, vinculin-positive focal adhesions, and intercellular vesicle transfer were previously described ^19,23^.

#### Scanning Electron Microscopy (SEM) and correlative light/SEM

CAD and SH-SY5Y cells were plated overnight on gridded IBIDI dishes, fixed at 37°C in 0.05% glutaraldehyde, 2% PFA in 0.2 M Hepes for 15 min followed by fixation in 4% PFA in 0.2 M Hepes for 15 min, stained with WGA 488 (1:300) for 15 min a RT. The samples were post-fixed in 2.5% glutaraldehyde in 0.2 M cacodylate buffer (pH 7.2) at 4°C, washed three times 5 mn in 0.2M cacodylate buffer (pH 7.2), treated for 1h with 1% osmium tetroxide in 0.2 M cacodylate buffer and then rinsed in distilled water. Samples were dehydrated through a graded series of 25, 50, 75 and 95% ethanol solutions for 5 min and for 10 min in 100% ethanol followed by critical point drying with CO2. Samples were sputtered with a 10 nm gold/palladium layer and were observed in a JEOL JSM-6700F field emission scanning electron microscope at a voltage of 5kV.

#### Focused-ion Beam Scanning Electron Microscopy (FIB-SEM)

SH-SY5Y cells were plated on gridded Ibidi μ-dishes (Biovalley, France). Positions were identified and recorded by light and fluorescence microscopy. After chemical fixation (same as described for SEM and correlative light/SEM), cells were incubated in 1% (w/v) osmic acid and 1.5 % (w/v) potassium ferrocyanide for 30 min, 1% tannic acid for 30 min, and 1% osmic acid for 30 min. Samples were dehydrated in a graded ethanol series (25%, 50%, 75%, 95% and 100%) and embedded in Epon (Agar Scientific, UK) and subsequently placed on a pin stub and recovered with silver paint (Agar Scientific, UK). Flat surfaces were coated with a 10 nm-thick layer of gold/palladium using an ion Beam Coater (Gatan Inc, USA) to avoid charging effect. 3D tomography was made with a FIB-SEM Auriga (Zeiss, Germany). An additional 1μm protective layer of platinum was deposited on the surface of the region of interest using an ion beam assisted deposition with 30 kV acceleration potential. The cross-section was milled using a 10 nA ion beam current. The surface obtained was then polished using a 2 nA ion beam current. Tomographic sections of 10 nm during acquisition were obtained using a 500 μA ion beam current. SEM images were acquired at 10 nm per pixel for a frame size window of 2048*2048 pixels by using a 1.5 keV acceleration voltage and a 30 μm aperture with the back-scattered electrons detector. Alignment of the acquired stack of images was done using ImageJ. Segmentations and measurements were performed using Amira 6.4 (Thermo Fisher Scientific, USA).

#### Cryo-correlative Light and Electron Microscopy (i-CLEM)

Vitrified TEM grids containing fluorescently labelled CAD cells were imaged on an epifluorescent Axiovert 200M inverted microscope (Zeiss) equipped with a cryo-correlative stage (Cryostage2, FEI), 10x (NA 0.3), 40x (NA 0.6) and 63x (NA 0.75, working distance 1.7 mm) long working distance air objectives and with a 482/18 (Excitation), 520/28 (Emission, green) filter cube. Fluorescent and phase contrast imaging at cryogenic temperatures was performed as previously described ^45−48^. Briefly, the cryo-correlative stage (MPI Biochemistry, Martinsried ^47^ was mounted onto the light microscope motorized stage and samples were transferred from liquid nitrogen storage only when the temperature of the microscope cryo-correlative stage was below −170 °C. Digital images were recorded with a AxiocamMRm camera (Zeiss). Following cryo-fluorescence imaging, the vitrified samples on grids were stored in liquid nitrogen until they were used for cryo-TEM. In all images, the brightness and contrast were adjusted in order to highlight TNTs.

#### Correlative Light and Cryo-electron Microscopy (ii-CLEM)

CAD cells on TEM finder grids were chemically fixed after fluorescent labeling as described in the cell culture section. The samples on TEM grids were then positioned in glass-bottom dishes (MatTek Corporation, Ashland, USA) in Hepes buffer. Fluorescence and corresponding transmitted light z-stack images of cells were obtained by using the same epifluorescence Axiovert 200M inverted microscope (Zeiss) equipped with 482/18 (Excitation), 520/28 (Emission, green) and a 546/12 (Excitation), 575-640nm (Emission, red) filter cubes and a 40x (NA 1.3, WD 0,21 mm) oil objective. Following fluorescence imaging, the samples on grids were immediately vitrified and stored in liquid nitrogen until they were used for cryo-TEM.

#### Cryo-electron Microscopy

Cryo-electron tomography was performed on a Tecnai 20 equipped with a field emission gun and operated at 200 kV (Thermo Fisher company). Images were recorded using either Explore 3D or SerialEM software on a 4k x 4k camera (Ultrascan from Gatan) and a Falcon II (FEI, Thermo Fisher) direct electron detector, with a 14μm pixel size. Tilt series of TNTs were acquired covering either an angular range of −52° to +52° (Supplementary Video 5), −51° to 51° (Fig. 4I and Supplementary Video 4), or in specific cases due to sample physical constraints with a reduced angular range (Fig. 4D, 4E, 4F, and corresponding Supplementary Videos 1, 2, and 3), with 2-4 degrees increment. The defocuses used were −8 and −10 μm. Tilt series of filopodial protrusions were acquired covering a tilt range of −65° to +66° (Fig. 6B, 6D, 6F, with corresponding Supplementary Videos 6, 7, 8, and Fig. 6H with corresponding Supplementary Video 9), with 2 degrees increment, defocuses of −6 μm. All tilt series were acquired at magnifications of 25,000x or 29,000x, binning 2, corresponding to a pixel size of 0.742 nm or 0,634 nm, respectively.

#### Live-Series Microscopy

Live time series images were acquired with a 60x 1.4NA CSU oil immersion objective lens on an inverted Elipse Ti microscope system (Nikon Instruments, Melville, NY, USA). Laser illumination was provided by 488 nm and 561 nm. Pairs of images were captured in immediate succession with one of two cooled CCD cameras, which enabled time intervals between 20 and 30 sec per z-stack. For live cell imaging, the 37 °C temperature was controlled with an Air Stream Stage Incubator which also controlled humidity. Cells were incubated with 5% CO2 during image acquisition.

### Image analysis and visualization

Tomographic tilt series were processed using version 4.9.2 of IMOD ^49^. Projections were pre-processed by hot pixel removal and rough alignment by cross-correlation. Final alignments were done by using 10 nm fiducial gold particles coupled to protein A gold. The reconstructions were obtained by using a weighted back-projection algorithm ^50^. For visualization purposes, the reconstructed volumes were processed by a Gaussian filter. Surface rendering was done with Amira 6.2 software package. Quantitative measurements of TNTs’ and vesicle diameter on TEM micrographs or on 3D tomographic reconstructions were done by using the open source program Fiji.

### Statistical analysis

The results of image analysis from pharmacological assays were transferred to Prism (GraphPad). For more than two groups statistical significance was assessed by a one-way ANOVA with Tukey correction. Differences were considered significant at \**p*< 0.05, \*\**p*< 0.005, or \*\*\**p*< 0.0005. Quantifications were done blind. Quantitative data depicted as (±SEM) mean standard deviation.

## Acknowledgements

We thank Seng Zhu for image analysis technical expertise and Christel Brou for valuable manuscript comments. We also gratefully acknowledge Gerard Péhau-Arnaudet (Ultrapole, Institut Pasteur) for his help in setting up freezing conditions of CAD cells during early stages of the project, and Remi Blanc from Amira (FEI, Thermo Fischer Scientific) for the automated segmentation of actin in Fig. 6 using the XTracing - Filament detection module. We thank the equipment excellence CACSICE for providing the Falcon II direct electron detector. We acknowledge the Ultrastructural Bio-Imaging facility at Institut Pasteur, member of the national infrastructure France-BioImaging (FBI) supported by the French National Research Agency (ANR-10-INBS-04). This work was supported by grants from the Agence Nationale de Recherche (JPND Neutargets: ANR-14-JPCD-0002-01 and ANR-16 CE160019-01 NEUROTUNN), and the Equipe FRM (Fondation Recherche Médicale) 2014 (DEQ20140329557) to C.Z. D.C.C was supported by the Pasteur - Paris University (PPU) International PhD Program. A.P. was supported by fellowships from France Parkinson and the Fondazione Ermenegildo Zegna.

## Author Contributions

A.S. performed all correlative, cryo-correlative light, and electron microscopy experiments and quantification, wrote the results and prepared the figures. D.C.C. prepared cells for TEM experiments, performed FM experiments and live imaging, helped with acquisition, prepared figures and wrote the manuscript. A.P. performed and analyzed CK-666 pharmacological experiments with D.C.C., set experimental conditions for SH-SY5Y cells, performed experiments and helped with the image rendering of TEM tomograms. K.G. and E.D. set up and prepared CAD cell cultures for TEM and SEM and discussed earlier experiments. S. C.-D. performed FIB-SEM experiments and the corresponding data rendering, C.S. performed SEM experiments in CAD cells. C.Z. conceived the project, supervised the work, and wrote the manuscript. A.S., D.C.C., A.P., J.K.-L., and C.Z. discussed the results. All authors commented on the manuscript.

## References

1 Rustom, A., Saffrich, R., Markovic, I., Walther, P. & Gerdes, H. H. Nanotubular highways for intercellular organelle transport. Science 303, 1007–1010, doi:10.1126/science.1093133 (2004).

2 Abounit, S. & Zurzolo, C. Wiring through tunneling nanotubes--from electrical signals to organelle transfer. Journal of cell science 125, 1089–1098, doi:10.1242/jcs.083279 (2012).

3 Austefjord, M. W., Gerdes, H. H. & Wang, X. Tunneling nanotubes: Diversity in morphology and structure. Communicative & integrative biology 7, e27934, doi:10.4161/cib.27934 (2014).

4 Vignais, M. L., Caicedo, A., Brondello, J. M. & Jorgensen, C. Cell Connections by Tunneling Nanotubes: Effects of Mitochondrial Trafficking on Target Cell Metabolism, Homeostasis, and Response to Therapy. Stem cells international 2017, 6917941, doi:10.1155/2017/6917941 (2017).

5 Ariazi, J. et al. Tunneling Nanotubes and Gap Junctions-Their Role in Long-Range Intercellular Communication during Development, Health, and Disease Conditions. Frontiers in molecular neuroscience 10, 333, doi:10.3389/fnmol.2017.00333 (2017).

6 Koyanagi, M., Brandes, R. P., Haendeler, J., Zeiher, A. M. & Dimmeler, S. Cell-to-cell connection of endothelial progenitor cells with cardiac myocytes by nanotubes: a novel mechanism for cell fate changes? Circulation research 96, 1039–1041, doi:10.1161/01.RES.0000168650.23479.0c (2005).

7 Abounit, S. et al. Tunneling nanotubes spread fibrillar alpha-synuclein by intercellular trafficking of lysosomes. The EMBO journal 35, 2120–2138, doi:10.15252/embj.201593411 (2016).

8 Abounit, S., Wu, J. W., Duff, K., Victoria, G. S. & Zurzolo, C. Tunneling nanotubes: A possible highway in the spreading of tau and other prion-like proteins in neurodegenerative diseases. Prion 10, 344–351, doi:10.1080/19336896.2016.1223003 (2016).

9 Chinnery, H. R., Pearlman, E. & McMenamin, P. G. Cutting edge: Membrane nanotubes in vivo: a feature of MHC class II+ cells in the mouse cornea. Journal of immunology 180, 5779–5783 (2008).

10 Lou, E. et al. Tunneling nanotubes provide a unique conduit for intercellular transfer of cellular contents in human malignant pleural mesothelioma. PloS one 7, e33093, doi:10.1371/journal.pone.0033093 (2012).

11 Osswald, M. et al. Brain tumour cells interconnect to a functional and resistant network. Nature 528, 93–98, doi:10.1038/nature16071 (2015).

12 Baker, M. How the Internet of cells has biologists buzzing. Nature 549, 322–324, doi:10.1038/549322a (2017).

13 Mattila, P. K. & Lappalainen, P. Filopodia: molecular architecture and cellular functions. Nature reviews. Molecular cell biology 9, 446–454, doi:10.1038/nrm2406 (2008).

14 Gutierrez-Vazquez, C., Villarroya-Beltri, C., Mittelbrunn, M. & Sanchez-Madrid, F. Transfer of extracellular vesicles during immune cell-cell interactions. Immunological reviews 251, 125–142, doi:10.1111/imr.12013 (2013).

15 Sowinski, S. et al. Membrane nanotubes physically connect T cells over long distances presenting a novel route for HIV-1 transmission. Nature cell biology 10, 211–219, doi:10.1038/ncb1682 (2008).

16 Kumar, A. et al. Influenza virus exploits tunneling nanotubes for cell-to-cell spread. Scientific reports 7, 40360, doi:10.1038/srep40360 (2017).

17 Lu, J. et al. Tunneling nanotubes promote intercellular mitochondria transfer followed by increased invasiveness in bladder cancer cells. Oncotarget 8, 15539–15552, doi:10.18632/oncotarget.14695 (2017).

18 Okafo, G., Prevedel, L. & Eugenin, E. Tunneling nanotubes (TNT) mediate long-range gap junctional communication: Implications for HIV cell to cell spread. Scientific reports 7, 16660, doi:10.1038/s41598-017-16600-1 (2017).

19 Delage, E. et al. Differential identity of Filopodia and Tunneling Nanotubes revealed by the opposite functions of actin regulatory complexes. Scientific reports 6, 39632, doi:10.1038/srep39632 (2016).

20 Gallo, G. Mechanisms underlying the initiation and dynamics of neuronal filopodia: from neurite formation to synaptogenesis. International review of cell and molecular biology 301, 95–156, doi:10.1016/B978-0-12-407704-1.00003-8 (2013).

21 Jacquemet, G., Hamidi, H. & Ivaska, J. Filopodia in cell adhesion, 3D migration and cancer cell invasion. Current opinion in cell biology 36, 23–31, doi:10.1016/j.ceb.2015.06.007 (2015).

22 Gousset, K. et al. Prions hijack tunnelling nanotubes for intercellular spread. Nature cell biology 11, 328–336, doi:10.1038/ncb1841 (2009).

23 Abounit, S., Delage, E. & Zurzolo, C. Identification and Characterization of Tunneling Nanotubes for Intercellular Trafficking. Current protocols in cell biology 67, 12 10 11–21, doi:10.1002/0471143030.cb1210s67 (2015).

24 Smith, I. F., Shuai, J. & Parker, I. Active generation and propagation of Ca2+ signals within tunneling membrane nanotubes. Biophysical journal 100, L37–39, doi:10.1016/j.bpj.2011.03.007 (2011).

25 Dieriks, B. V. et al. alpha-synuclein transfer through tunneling nanotubes occurs in SH-SY5Y cells and primary brain pericytes from Parkinson’s disease patients. Scientific reports 7, 42984, doi:10.1038/srep42984 (2017).

26 Schultz, M. Rudolf Virchow. Emerging Infectious Diseases 14, 1480–1481, doi:10.3201/eid1409.086672 (2008).

27 Gousset, K., Marzo, L., Commere, P. H. & Zurzolo, C. Myo10 is a key regulator of TNT formation in neuronal cells. Journal of cell science 126, 4424–4435, doi:10.1242/jcs.129239 (2013).

28 Patla, I. et al. Dissecting the molecular architecture of integrin adhesion sites by cryo-electron tomography. Nature cell biology 12, 909–915, doi:10.1038/ncb2095 (2010).

29 Korobova, F. & Svitkina, T. Arp2/3 complex is important for filopodia formation, growth cone motility, and neuritogenesis in neuronal cells. Molecular biology of the cell 19, 1561–1574, doi:10.1091/mbc.E07-09-0964 (2008).

30 Barzik, M., McClain, L. M., Gupton, S. L. & Gertler, F. B. Ena/VASP regulates mDia2-initiated filopodial length, dynamics, and function. Molecular biology of the cell 25, 2604–2619, doi:10.1091/mbc.E14-02-0712 (2014).

31 Medalia, O. et al. Organization of actin networks in intact filopodia. Current biology: CB 17, 79–84, doi:10.1016/j.cub.2006.11.022 (2007).

32 Aramaki, S., Mayanagi, K., Jin, M., Aoyama, K. & Yasunaga, T. Filopodia formation by crosslinking of F-actin with fascin in two different binding manners. Cytoskeleton 73, 365–374, doi:10.1002/cm.21309 (2016).

33 Pasquier, J. et al. Preferential transfer of mitochondria from endothelial to cancer cells through tunneling nanotubes modulates chemoresistance. Journal of translational medicine 11, 94, doi:10.1186/1479-5876-11-94 (2013).

34 Ady, J. *W.* et al. Intercellular communication in malignant pleural mesothelioma: properties of tunneling nanotubes. Frontiers in physiology 5, 400, doi:10.3389/fphys.2014.00400 (2014).

35 Zhu, H. et al. Rab8a/Rab11a regulate intercellular communications between neural cells via tunneling nanotubes. Cell death & disease 7, e2523, doi:10.1038/cddis.2016.441 (2016).

36 Wang, X. & Gerdes, H. H. Transfer of mitochondria via tunneling nanotubes rescues apoptotic PC12 cells. Cell death and differentiation 22, 1181–1191, doi:10.1038/cdd.2014.211 (2015).

37 Keller, K. E., Bradley, J. M., Sun, Y. Y., Yang, Y. F. & Acott, T. S. Tunneling Nanotubes are Novel Cellular Structures That Communicate Signals Between Trabecular Meshwork Cells. Investigative ophthalmology & visual science 58, 5298–5307, doi:10.1167/iovs.17-22732 (2017).

38 Gurke, S., Barroso, J. F. & Gerdes, H. H. The art of cellular communication: tunneling nanotubes bridge the divide. Histochemistry and cell biology 129, 539–550, doi:10.1007/s00418-008-0412-0 (2008).

39 Onfelt, B. et al. Structurally distinct membrane nanotubes between human macrophages support long-distance vesicular traffic or surfing of bacteria. Journal of immunology 177, 8476–8483 (2006).

40 Melkov, A. & Abdu, U. Regulation of long-distance transport of mitochondria along microtubules. Cellular and molecular life sciences: CMLS 75, 163–176, doi:10.1007/s00018-017-2590-1 (2018).

41 Morris, R. L. & Hollenbeck, P. J. Axonal transport of mitochondria along microtubules and F-actin in living vertebrate neurons. The Journal of cell biology 131, 1315–1326 (1995).

42 Quintero, O. A. et al. Human Myo19 is a novel myosin that associates with mitochondria. Current biology: CB 19, 2008–2013, doi:10.1016/j.cub.2009.10.026 (2009).

43 Pathak, D., Sepp, K. J. & Hollenbeck, P. J. Evidence that myosin activity opposes microtubule-based axonal transport of mitochondria. The Journal of neuroscience: the official journal of the Society for Neuroscience 30, 8984–8992, doi:10.1523/JNEUROSCI.1621-10.2010 (2010).

44 Dubochet, J. et al. Cryo-electron microscopy of vitrified specimens. Quarterly reviews of biophysics 21, 129–228 (1988).

45 Sartori, A. et al. Correlative microscopy: Bridging the gap between fluorescence light microscopy and cryo-electron tomography. J Struct Biol 160, 135–145, doi:10.1016/j.jsb.2007.07.011 (2007).

46 Lucic, V. et al. Multiscale imaging of neurons grown in culture: from light microscopy to cryo-electron tomography. J Struct Biol 160, 146–156, doi:10.1016/j.jsb.2007.08.014 (2007).

47 Rigort, A. et al. Micromachining tools and correlative approaches for cellular cryo-electron tomography. J Struct Biol 172, 169–179, doi:10.1016/j.jsb.2010.02.011 (2010).

48 Hampton, C. M. et al. Correlated fluorescence microscopy and cryo-electron tomography of virus-infected or transfected mammalian cells. Nature protocols 12, 150–167, doi:10.1038/nprot.2016.168 (2017).

49 Kremer, J. R., Mastronarde, D. N. & McIntosh, J. R. Computer visualization of three-dimensional image data using IMOD. J Struct Biol 116, 71–76, doi:10.1006/jsbi.1996.0013 (1996).

50 Hegerl, R. The EM Program Package: A Platform for Image Processing in Biological Electron Microscopy. J Struct Biol 116, 30–34 (1996).

